# microRNAs bidirectionally regulate FUT1 to modulate α-1,2-fucosylation and cancer-associated biology

**DOI:** 10.64898/2026.03.13.711703

**Authors:** Tigist Batu, Chu Thu, Lara K. Mahal

## Abstract

The α-1,2-fucosyltransferase, FUT1, plays a central role in blood type determination, the establishment of the gut microbiota, and cancer progression. Using high-throughput analysis, we mapped the miRNA regulatory landscape of FUT1 and found that miRNAs bidirectionally regulate this enzyme. Validation of miRFluR assay results across multiple cell lines confirmed that both upregulatory and downregulatory miRNA interactions affect endogenous FUT1 and its enzymatic product, α-1,2-fucosylation. Inhibitors of endogenous miRNA impacted both enzyme and glycan levels, underscoring the biological impact of bidirectional miRNA regulation. Upregulatory binding sites (miR-200c-5p and miR-361-3p) and the downregulatory binding site (miR-29c-5p) identified in this work both displayed non-canonical binding. Notably, miRNAs upregulating FUT1 are depleted in various cancers, aligning with observations that loss of α-1,2-fucosylation is a hallmark of esophageal cancer, melanoma biogenesis, and metastasis. Furthermore, integration of our previous work with the current results indicates that the loss of miR-200c family has a strong correlation with the loss of α-1,2-fucosylation during epithelial-to-mesenchymal transition. Together, these results identify miRNAs as key regulators of FUT1 and demonstrate that bidirectional miRNA control of glycosyltransferases can reshape cell-surface glycosylation with important implications for cancer biology.

## Introduction

Fucosylation is a critical glycan modification that influences cell-cell communication, signal transduction, and host–microbe interactions. Among the enzymes catalyzing this process, α-1,2-fucosyltransferase-1 (FUT1) plays a pivotal role by transferring fucose residues in an α-1,2 linkage to galactose, generating the H-antigen that serves as a precursor for ABO blood groups, as well as Lewis antigens such as Lewis y and Lewis b (1–3). Beyond its well-established role in hematology and immunology, changes in FUT1-mediated α-1,2-fucosylation have increasingly been implicated in pathological processes, including cancer progression, inflammation, and epithelial-to-mesenchymal transition (EMT) (4–7).

Aberrant expression of FUT1 has been reported across multiple cancers. Studies in colorectal, melanoma, and certain gastrointestinal cancers suggest that α-1,2-fucosylation may exert tumor-suppressive effects (7–9). Conversely, elevated FUT1 and the consequent accumulation of α-1,2-fucose has been associated with enhanced cell proliferation, invasion, metastasis, and chemoresistance in ovarian and breast cancer (5,6). These findings indicate the dual nature of FUT1 function and highlight the importance of understanding the mechanisms that regulate its expression.

MicroRNAs (miRNAs), small noncoding RNAs that post-transcriptionally regulate gene expression, have emerged as central players in controlling glycosylation enzymes (10–12). Although miRNAs are commonly known to downregulate protein expression, it is now evident that they can also have an upregulatory role. Multiple works from our laboratory and others have identified upregulation by miRNA in a variety of contexts (13–21). Recently, it has become clear, through high-throughput analysis of miRNA regulation, that bidirectional control by miRNA is the common state in dividing cells (16–21). These experiments have expanded our understanding of miRNA regulation.

Despite the recognized importance of FUT1 in cancer biology, the regulation of FUT1 expression at the post-transcriptional level remains poorly defined. A limited number of studies have hinted at miRNA involvement in the regulation of α-1,2-fucosylation, but a systematic analysis of miRNA:FUT1 interactions has not been done. Herein, we investigate the role of miRNAs in the regulation of FUT1 using our high-throughput miRFluR assay (16). We define the miRNA:FUT1 regulatory landscape, characterizing the downstream effects of FUT1 modulation by miRNA. By examining the miRNA regulatory map of FUT1, we identify new potential roles for this enzyme and α-1,2-fucosylation in cancer and infectious disease.

## Results

### miRFluR high-thoughput assay reveals FUT1 is predominantly upregulated by miRNA

To study the miRNAome regulation of FUT1, we utilized the high-throughput miRFluR assay developed in our lab (16) (**Fig. 2A**). miRFluR is a dual fluorescence assay system that utilizes a plasmid (pFmiR) in which the 3’UTR of a gene of interest, in this case FUT1, is cloned down-stream of the fluorescent protein Cerulean. A second fluorescent protein, mCherry, is present in the same vector under an identical promoter as an internal control (16–20). When cells are co-transfected with miRNA mimics, the ratio of Cerulean (Cer) to mCherry (mCh) indicates the impact of the miRNA on protein levels (e.g. downregulation: low Cer:mCh, upregulation: high Cer:mCh, **Fig. 2A**). The 3’UTR of FUT1 (ENST00000310160.3, 2173 nucleotides) is conserved in all isoforms of the transcript and was cloned into the pFmiR plasmid to make our pFmiR-FUT1 sensor (**Figs. S1 & S2**). We co-transfected HEK293T cells with pFmiR-FUT1 and a miRNA mimic library (Dharmacon, v. 21, 2601 miRNA) arrayed in triplicate in 384-well plates. After 72 hours (h) post-transfection, the fluorescence signal of the reporter proteins was recorded, and the Cer:mCh ratio was calculated. Although our plates contained non-targeting control (NTC), we found that for many plates the error in the NTC was above our exclusion threshold (>15%), thus we normalized the Cer:mCh ratio for each miRNA to the median Cer:mCh ratio for each plate. As a quality control (QC) measure, data points with greater than 15% measurement error or plates where the median error was > 15% were excluded from the dataset. Only 985 of the 2601 miRNAs passed our rigorous QC, which is a lower number than has been observed for other genes (16–20). In general, after optimization we observed lower levels of transfection with the pFmiR-FUT1 plasmid, causing the mCherry signals to be closer to the signal-to-noise threshold. This added noise to the dataset, resulting in fewer miRNAs meeting our criteria for inclusion. We z-scored our QCed data, considering hits as those within the 95% confidence interval, and found 46 miRNAs within our dataset that regulated FUT1, corresponding to ∼5% of the QCed data. (**Fig. 2B**). This is similar to other datasets obtained with our miRFluR assay (∼3-6% hit rate) (16–21). Of the 46 miRNA hits, 80% were upregulatory (norm. Cer/mCh >1.50, 39 upmiRs) and 20% were downregulatory (norm Cer/mCh < 0.57, 7 downmiRs) (**Fig. 2B**, **Dataset S1)**.

### FUT1 is bidirectionally regulated by miRNA in multiple cancer cell lines

In our previous collaborative work, we identified a loss of α-1,2-fucosylation upon transformation from nevi to primary to metastatic melanoma (8, 22). The loss of FUT1 mRNA expression correlated with melanoma progression in multiple datasets and silencing FUT1 led to increased melanoma cell invasion, in line with a role in metastasis (8). In one of our earliest miRNA papers, we identified a correlation between α-1,2-fucosylation and an epithelial phenotype, and found that the lung adenocarcinoma line A549 had intermediate levels of α-1,2-fucosylation, while the epithelial colon adenocarcinoma cell line HT29 displayed high levels of the epitope (23). Based on our previous studies, we chose the metastatic melanoma line A375, along with A549 and HT29 to validate our miRFluR results. We silenced of FUT1 with pooled siRNA in both A375 and HT29 to confirm the specificity of our antibody (**Figs. S3 & S4**). In previous work, we found that the Dharmacon non-targeting control (NTC) could impact protein levels through the 3’UTR in unexpected ways (18). Thus, we first tested the impact of NTC on FUT1 protein levels and found that the Dharmacon NTC upregulated FUT1 when compared to untreated cell lysate **(Fig. S4)**. Therefore, we tested two miRNAs from the middle of our dataset (miR-8066, −513b-3p, **Dataset S1**) for their impact on FUT1. These miRNA had few biological associations and no impact on FUT1 protein levels **(Figs. S4)**. We thus decided to use miR-8066 as our new NTC (NTC*) for all miRNA mimic experiments.

To validate our miRFluR dataset, we selected 5 upmiRs and 5 downmiRs from our hit list, focusing on miRNA that were well characterized, up-miRs (blue): miR-200c-5p, - 361-3p, −340-5p, −29b-3p, −143-5p; down-miRs (red): miR-29c-5p, −769-3p, −4524, −5047, −4757-5p) (**Fig. 2B**) (28). We transfected cells (A375, A549, HT29) with miRNA mimics or NTC*, lysed after 48 h and analyzed FUT1 protein levels by Western blot or mRNA using RT-PCR (**Figs. 3 & S5**). In general, our data confirmed the results of our miRFluR assay, with each miRNA significantly impacting FUT1 levels in at least one of the three cell lines. We observed some cell dependency. For up-miRs, four of the five tested miRNAs were significant in all three cell lines, with miR-29b-3p showing mild but insignificant upregulation in A549. In contrast, only one (miR-29c-5p) of the five down-miRs tested was significant in all three cell lines. In the melanoma line A375, up-miRs enhanced FUT1 expression by ∼30% to 2-fold, while down-miRs caused a ∼50% decrease (**Figs. 3A-B, S5A-B**). We observed a similar trend in A549, with up-miRs enhancing expression by ∼20-60% and down-miRs resulting in a ∼20-40% reduction (**Figs. 3A & C, S5C-D**). The data in HT29 shows ∼40-80% upregulation of FUT1 expression by up-miRs, but down regulation is muted with miR-29c-5p, the strongest in this setting, giving ∼30% reduction of FUT1 expression (**Figs. 3A & D, S5E-F**). In line with our previous studies, the impact of miRNA on the mRNA level of FUT1, determined using RT-PCR, showed mixed results, with some changes concordant with and some discordant with the protein level data (**Figs. 3E-F**). Overall, this data validates our miRFluR assay results and shows that the impact of miRNA is cell line dependent, in line with other studies (16–21).

### miRNA bidirectionally regulates α1,2-fucosylation

To study the impact of miRNA regulating FUT1 on α1,2-fucosylation, we used a fluorophore-conjugated lectin, *Ulex europaeus* agglutinin I (UEA-I) that predominantly binds to the α-1,2-fucose epitope (**Fig. 1**) (21). We transfected A375 and HT29 cells with either up-miRs (miR-200c-5p, −361-3p), down-miRs (miR-29c-5p, 769-3p) or NTC*. After 48 h, cells were fixed and stained with UEA-I. In line with our expectations, the up-miRs led to higher levels of α-1,2-fucosylation while down-miRs inhibited the expression of this epitope (**Figs. 4A-B, & S7**).

**Figure 1.**
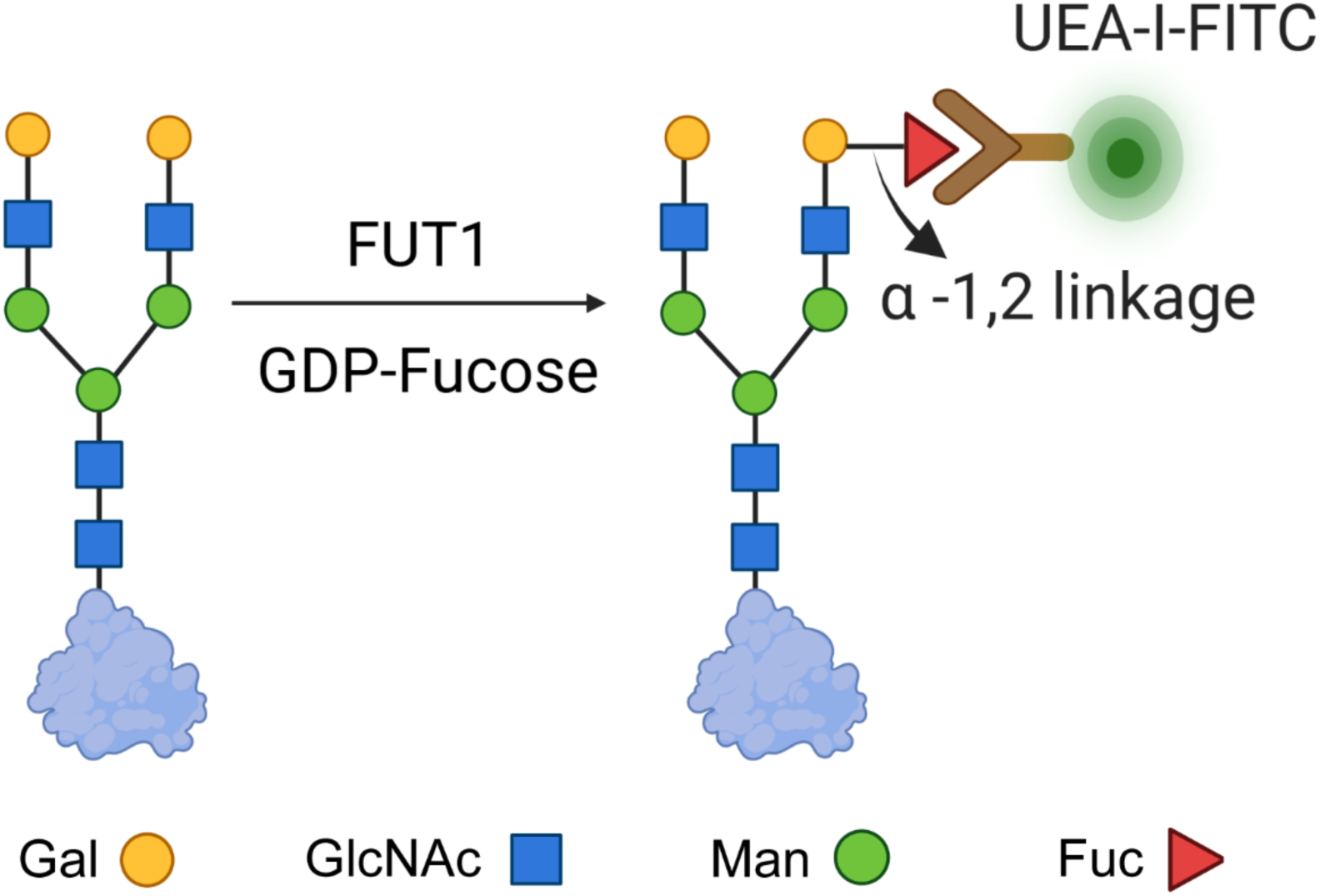
Scheme for α-1,2-Fucosylation. Enzymatic addition of Fucose to N- glycans by FUT1. SNFG symbols are shown (red triangle: fucose (Fuc), yellow circle: galactose (Gal), blue square: N-acetylglucosamine (GlcNAc)).

**Figure 2.**
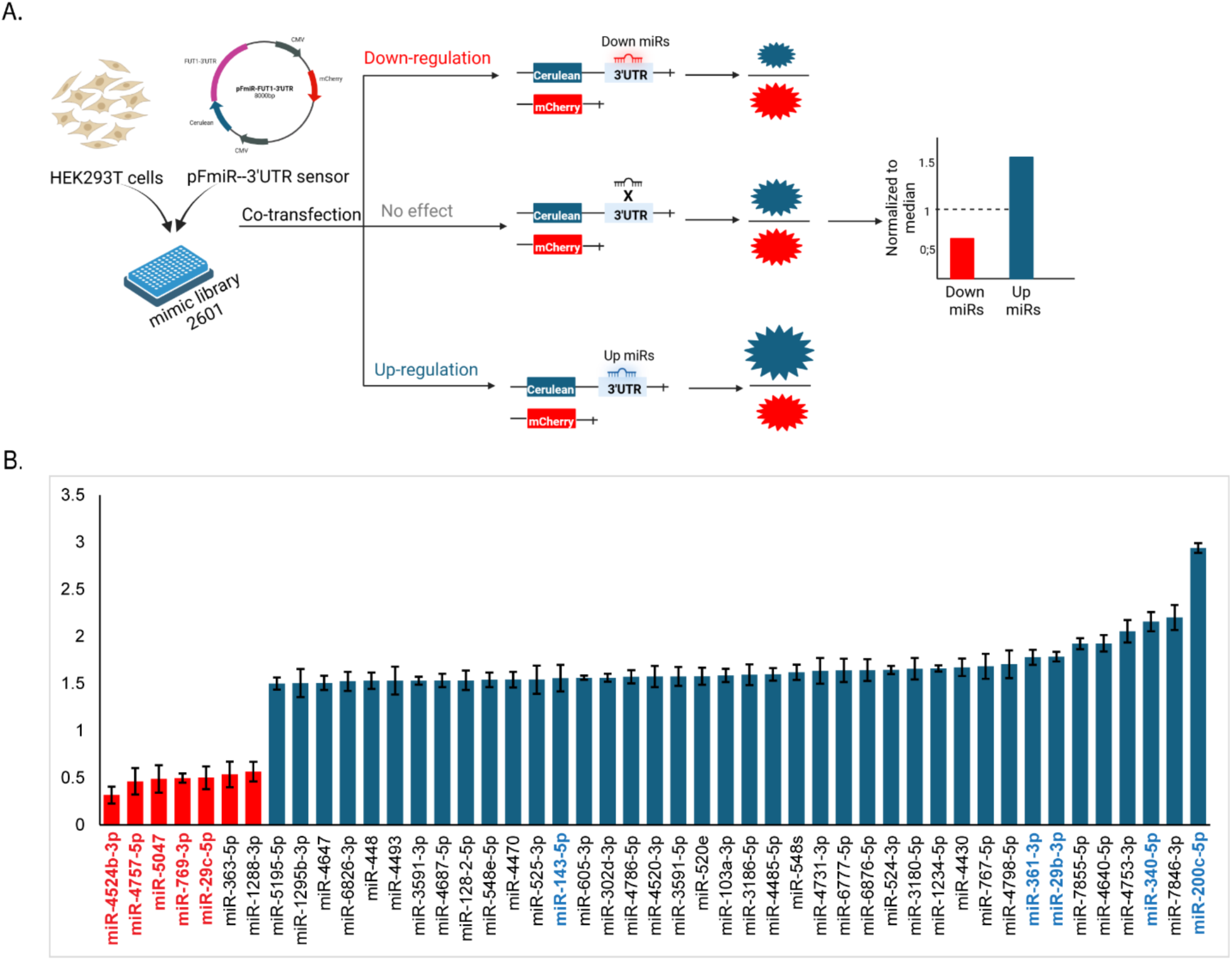
miRFluR high-throughput assay reveals FUT1 is predominantly upregulated by miRNA. (A) Schematic of the miRFluR assay. pFmiR-3’UTR FUT1 sensor is co-transfected with human miRNAome library and HEK293T cells in 384-well plates. (B) Bar graph of miRNA in the 95% confidence interval for FUT1 (down-miRs 20%: red, up-miRs 80%: blue). Data are normalized to the median. Error bars represent the error of measurement. miRNA in the 95% confidence interval for FUT1 regulation are colored (down-miRs: red, up-miRs: blue). All miRNA data shown are post-QC.

To more carefully quantify these results, we analyzed cells by flow cytometry (**Figs. 4C-H & S8**). Our results confirmed the lectin staining results in both A375 (**Fig. 4C-D**, ∼ 20% downregulation and ∼60% upregulation observed, respectively) and HT29 (**Fig. 4E-F**, ∼ 20% downregulation and ∼20% upregulation observed, respectively). We also tested the impact of these miRNAs in A549. Interestingly, in this cell line, while we observed the expected downregulation (**Fig. 4G-H**, ∼30-40%), the upregulators did not significantly impact α-1,2-fucosylation. This contrasts with the changes in FUT1 levels observed by Western blot (**Fig. 3A & C**). Overall, our data provide evidence that miRNA mediated regulation of FUT1 has a direct impact on α-1,2-fucosylation.

**Figure 3.**
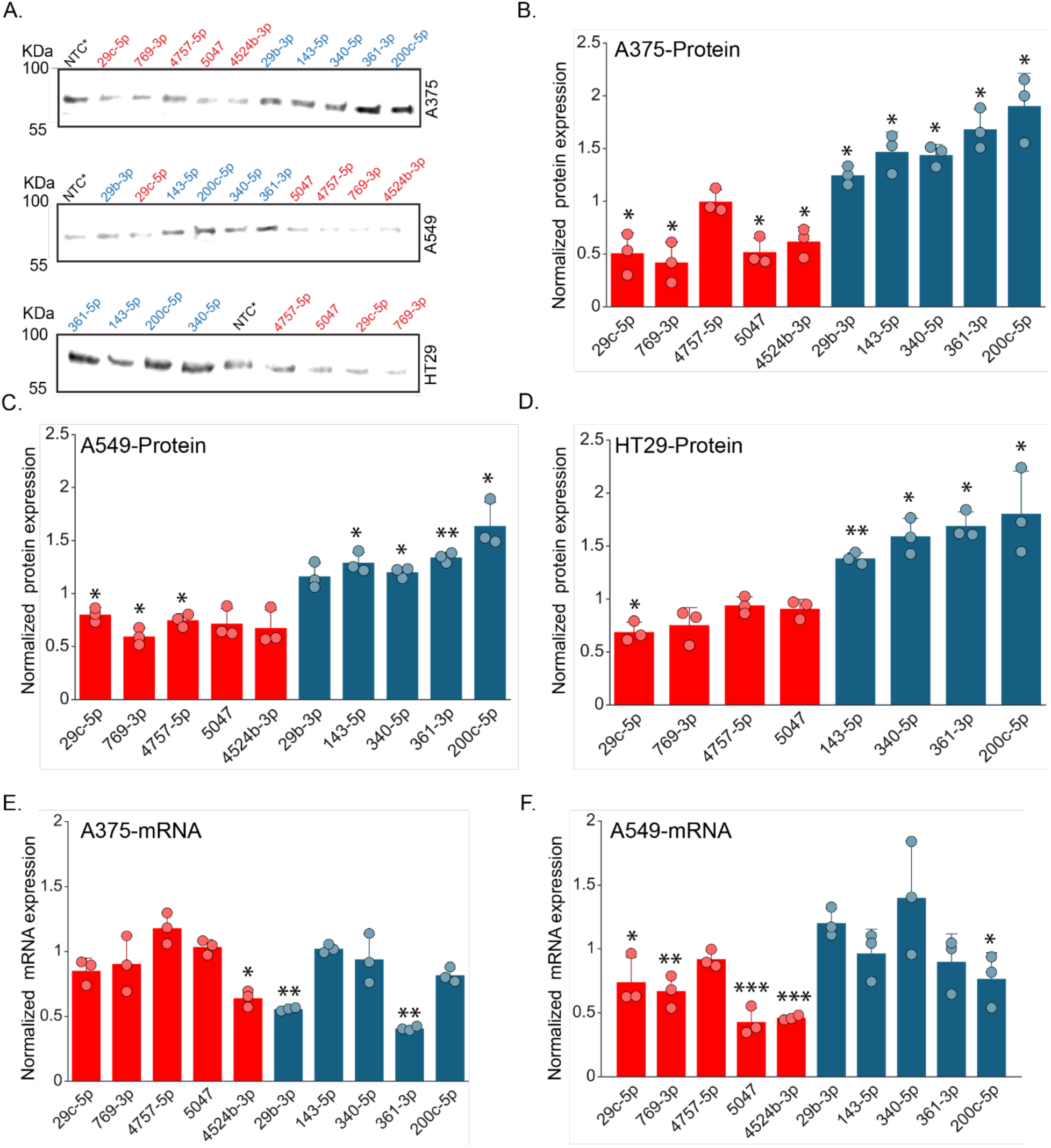
Validation of miRNA regulators of FUT1 in multiple cancer cell lines. (A) Representative western blot of showing FUT1 expression in A375 (top), A549 (middle), HT29 (bottom). (B) Quantification of Western blots of FUT1 in A375 cells. A375 cells were transfected with miRNA mimics or nontargeting control (NTC*, 50 nM, 48 h, n=3). (C) Quantification of Western blots of FUT1 in A549 cells. A549 cells were transfected with miRNA mimics or nontargeting control (NTC*, 50 nM, 48 h, n=3). (D) Quantification of Western blots of FUT1 in HT29 cells. HT29 cells were transfected with miRNA mimics or nontargeting control (NTC*, 50 nM, 48 h, n=3) (E) Quantification of RTqPCR result of FUT1 in A375. A375 cells were transfected with miRNA mimics or nontargeting control (NTC*, 50 nM, 48 h, n=3). Graph of normalized FUT1 mRNA expression relative to GAPDH. (F) Quantification of RTqPCR result of FUT1 in A549. A549 cells were transfected with miRNA mimics or nontargeting control (NTC*, 50 nM, 48 h, n=3). Graph of normalized FUT1 mRNA expression relative to GAPDH. For Western blots, normalized FUT1 signal for any miRNA was first normalized to Ponceau and then to the normalized NTC* signal. Dots indicate independent biological replicates. Errors shown are standard deviations. One-sample *t*-test was used to compare miRs to NTC (ns not significant, * *p* < 0.05, ** < 0.01, *** < 0.001). For whole blots and corresponding Ponceaus, see **Figure S4**.

**Figure 4.**
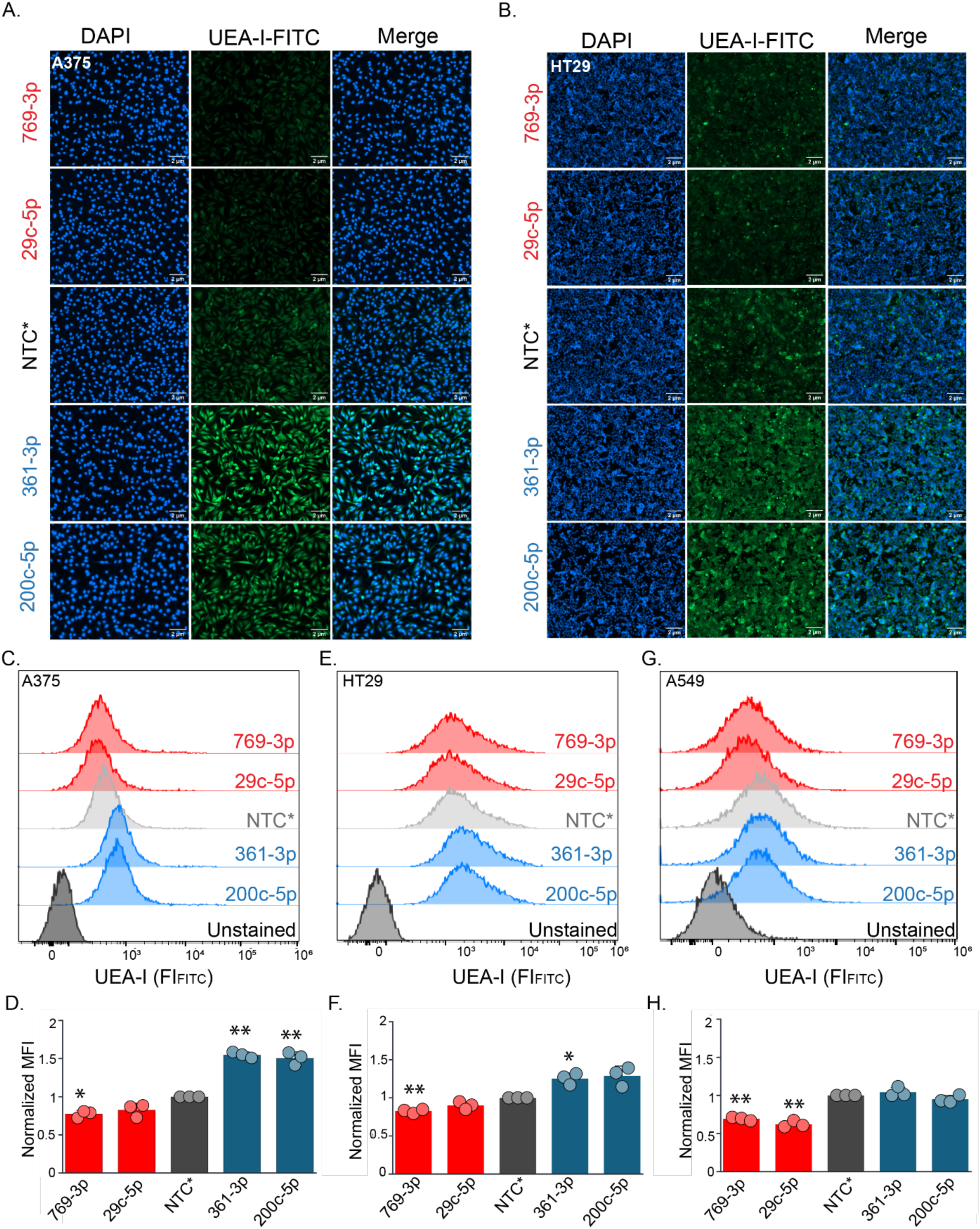
Validation of miRNA regulators of α-1,2-Fucose. (A) UEA-I staining of A375. A375 cells were treated as in Figure 3. After 48 hours, they were fixed with paraformaldehyde and stained with DAPI and FITC-UEA-I. (B). UEA-I staining of HT29. HT29 cells were treated as in Figure 3. After 48 hours, cells were fixed with paraformaldehyde and stained with DAPI and FITC-UEA-I. Images were taken after coverslips were mounted on a slide under fluorescence microscope slide. (C) Representative flow cytometry histogram for A375 cells (dark gray: unstained, gray: NTC*, blue: up-miRs, red: downmiRs). (D) Bar chart corresponding to C (MFI: mean fluorescence intensity). All experiments were performed in biological triplicate. (E)Representative flow cytometry histogram for HT29 cells (dark gray: unstained, gray: NTC*, blue: up-miRs, red: downmiRs). (F) Bar chart corresponding to E (MFI: mean fluorescence intensity). All experiments were performed in biological triplicate. (G)Representative flow cytometry histogram for A549 cells (dark gray: unstained, gray: NTC*, blue: up-miRs, red: downmiRs). (H) Bar chart corresponding to G (MFI: mean fluorescence intensity). All experiments were performed in biological triplicate. For flow cytometry quantification, dots indicate independent biological replicates. Errors shown are standard deviations. One-sample t-test was used to compare miRs to NTC* (* p < 0.05, ** < 0.01, *** < 0.001).

### Inhibition of endogenous miRNA confirms the impact of miRNA on FUT1

Anti-miRs are single-stranded oligonucleotides that are complementary to endogenous miRNAs, complexing with them in AGO2 and inhibiting their function (**Fig. 5A**) (25). The ability of anti-miRs to inhibit is dependent upon the endogenous expression levels of the miRNA. We tested the impact of anti-miRs for a subset of miRNAs in HT29 (anti-down-miRs: 29c-5p, 769-3p; anti-up-miRs: 200c-5p, 361-3p, 340-5p). In HT29, miRs-361-3p and −340-5p are highly expressed, miRs-29c-5p is moderately expressed, and low levels of expression of miR-200c-5p and −769-3p are observed (26). We transfected HT29 with anti-miRs or the supplied non-targeting control (anti-NTC, 50 nM) for 48 h and then performed Western blot analysis for FUT1 (**Figs. 5B, S6**). In general, our data are in line with expectations, with anti-down-miRs showing an increase in FUT1 protein levels (∼10-20%) and the anti-up-miRs showing a decrease (20-30%). However, no inhbition was observed for anti-200c-5p, which is poorly expressed in this cell line. In line with our findings, inhibiting endogenous miRNA impacted α-1,2-fucosylation levels, with anti-downmiR-29c-5p increasing, and anti-upmiR-361-3p decreasing, expression of the epitope **(Fig. 5C-D)**. Overall, our data showcases the impact of endogenous miRNA on FUT1 expression and α-1,2-fucosylation.

**Figure 5.**
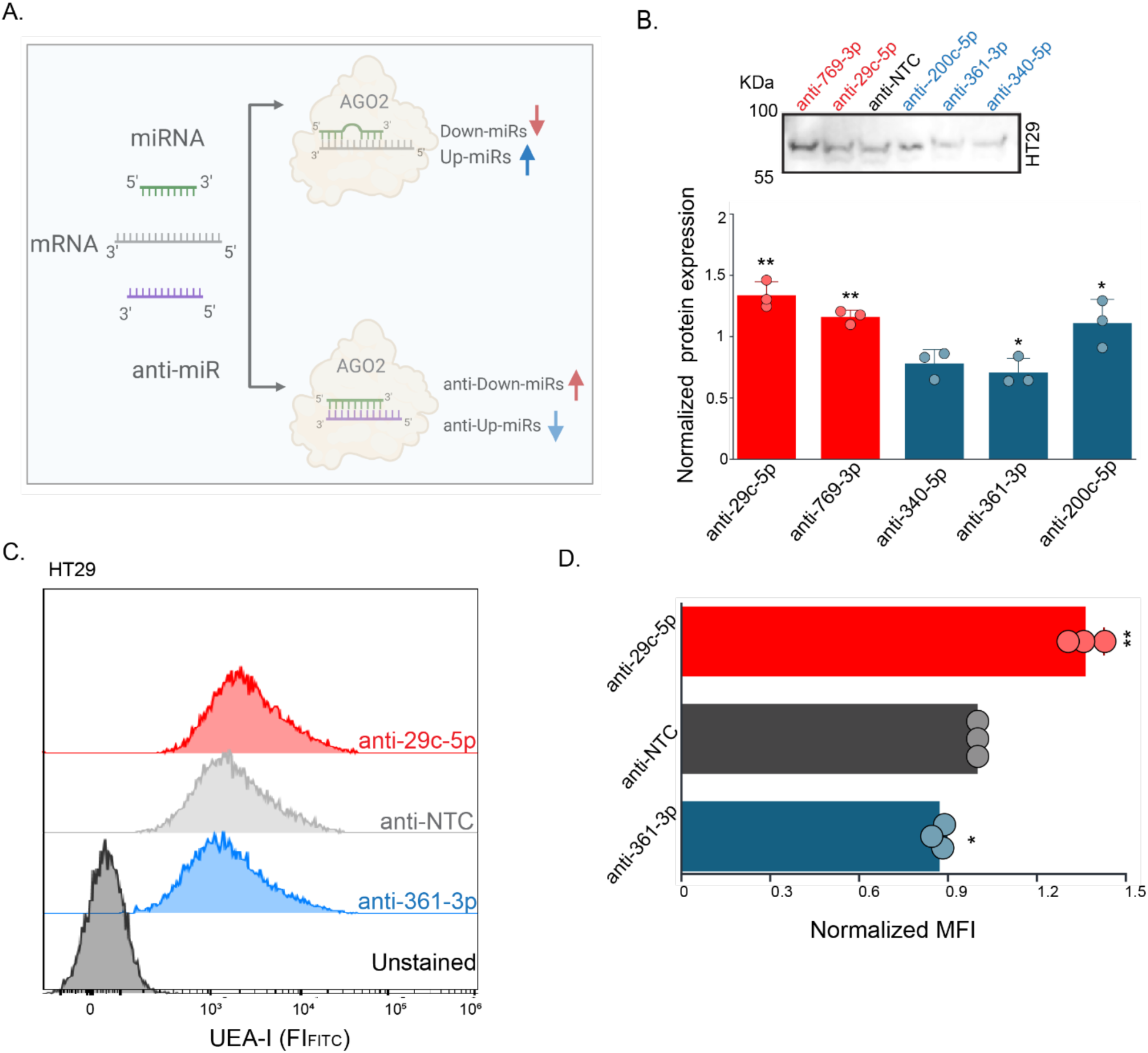
Inhibition of endogenous miRNAs impacts FUT1 and α-1,2-Fucose. (A) Schematic representation of anti-miR function. (B) Quantification of Western blots of FUT1 (n=3). HT29 cells were transfected with anti-miR or matched nontargeting control (anti-NTC, 50 nM, 48 h). The inset shows a representative blot. FUT1 expression was normalized by Ponceau and divided by the normalized signal from anti-NTC. (C-D) Flow cytometry analysis (n=3). All cells were treated with anti-miRs as in B and stained with FITC-UEA-I (D). Representative histogram (dark gray: unstained, gray: anti-NTC., blue: anti-upmiR, red: anti-downmiR). (E) Bar chart corresponding to D. Dots indicate independent biological replicates. Errors shown are standard deviations. One sample *t*-test was used to compare anti-miRs to anti-NTC ( * *p* < 0.05, ** < 0.01, *** < 0.001).

### miRNAs regulate FUT1 via direct interactions with the 3’UTR

In our previous work, we found that our miRFluR analysis identifies miRNA that directly interact with the 3’UTR to regulate the gene of interest (28–30). To identify sites for the down-miR (miR-29c-5p) and up-miRs (miR-200c-5p and −361-3p), we surveyed Targetscan. This algorithm predicts canonical binding sites, which have seed binding between nucleotide 2 and 8 in the miRNA (32). No predictions were found for any of these three miRNAs. We then surveyed RNAhybrid, a program that identifies the most stable binding sites between miRNA and mRNA (30). We found sites for all three miRNAs with minimum free energies of <-20 kcal/mol (**Fig. 6A-C**). All three sites were non-canonical, including that of the down-miR. We mutated these predicted binding sites in the pFmiR-FUT1 sensor to the respective miRNA sequence to generate a series of mutant sensors. These sensors were co-transfected with the appropriate miRNA mimics in HEK293T cells, and the fluorescence intensity was measured after 72 h. Data was normalized over the NTC* for each sensor (**Fig. 6D-F**). For all the three miRNAs (down-miR 29c-5p and up-miRs −200c-5p and −361-3p) the mutations significantly impacted miRNA regulation (p<0.05), validating that the impact of these miRNAs are via direct interactions.

**Figure 6.**
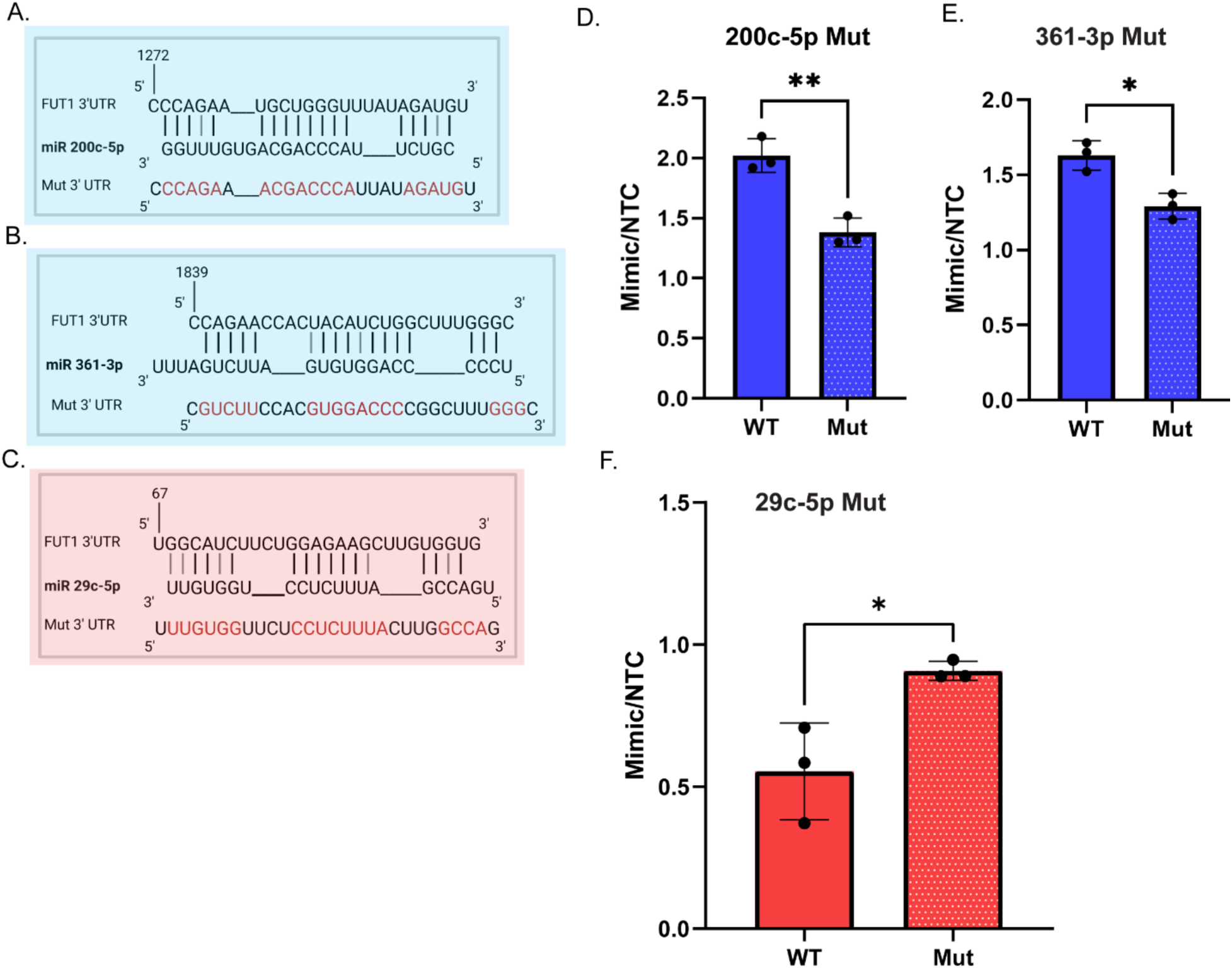
FUT1 miRNA regulation is via non-canonical interactions of miRs with the 3’UTR. (A) Alignment of miR-200c-5p with their predicted FUT1-3′UTR sites and the corresponding mutants. Mutated residues are shown in red, wobble interactions (G: U) are indicated in grey. (B) Alignment of miR-361-3p with their predicted FUT1-3′UTR sites and the corresponding mutants. Mutated residues are shown in red, wobble interactions (G: U) are indicated in grey. (C) Alignment of miR-29c-5p with their predicted FUT1-3′UTR sites and the corresponding mutants. Mutated residues are shown in red, wobble interactions (G: U) are indicated in grey. (D) Bar graph of data from wildtype (WT, solid) and mutant (200c-5p MUT, square) miRFluR sensors. (E) Bar graph of data from wildtype (WT, solid) and mutant (361-3p MUT, square) (F) Bar graph of data from wildtype (WT, solid) and mutant (29c-5p MUT, square). For each sensor, data was normalized over NTC*. The experiment was performed in biological triplicate, and the paired *t-*test was used for comparison. (* p < 0.05, ** < 0.01, *** < 0.001).

### miRNAs upregulating FUT1 are depleted in cancer and associate with EMT

We performed enrichment analysis for pathways and diseases associated with the up-miRs regulating FUT1 using miEAA, an online miRNA enrichment tool (**Table 1, Dataset 2**) (27). We only considered pathways and diseases in which at least 10% of the input list (4 miRs) were represented. Depletion of up-miRs for FUT1 was strongly associated with multiple cancers, including colon, lung, and melanoma. This is consistent with data from other papers in which the loss of α-1,2-fucosylation and FUT1 is associated with transformation and cancer progression (8,22,28).

**Table 1.**

| **Disease enrichment (MNDR)** |  |  |  |
| --- | --- | --- | --- |
| **Subcategory** | **Enrichment** | **P-value** | **# miRs** |
| *Colon adenocarcinoma* | depleted | 0.001 | 4 |
| *Head & neck squamous cell carcinoma* | depleted | 0.006 | 6 |
| *Burkitt lymphoma* | depleted | 0.011 | 5 |
| *Prostate cancer* | depleted | 0.025 | 7 |
| *Lung squamous cell carcinoma* | depleted | 0.028 | 10 |
| *Breast carcinoma* | depleted | 0.030 | 4 |
| *Pituitary adenoma* | depleted | 0.042 | 4 |
| *Carcinoma renal cell* | depleted | 0.044 | 6 |
| *Kidney chromophobe cancer* | depleted | 0.044 | 6 |
| *Lung small cell carcinoma* | depleted | 0.044 | 6 |
| *Small cell lung cancer* | depleted | 0.044 | 6 |
| **Pathway enrichment (KEGG)** |  |  |  |
| **Subcategory** | **Enrichment** | **P-value** | **# miRs** |
| Inositol phosphate metabolism | depleted | 0.003 | 18 |
| Proteasome | depleted | 0.003 | 8 |
| Pentose and glucuronate interconversions | depleted | 0.003 | 4 |
| Cell adhesion molecules | depleted | 0.003 | 20 |
| ABC transporters | depleted | 0.003 | 10 |
| Arachidonic acid metabolism | depleted | 0.003 | 10 |
| IL-17 signaling pathway | depleted | 0.003 | 21 |
| Neuroactive ligand-receptor interaction | enriched | 0.003 | 27 |
| cAMP signaling pathway | enriched | 0.003 | 31 |
| Non-homologous end-joining | depleted | 0.003 | 4 |
| Autophagy - animal | depleted | 0.003 | 22 |
| N-Glycan biosynthesis | depleted | 0.003 | 12 |
| Bacterial invasion of epithelial cells | enriched | 0.003 | 22 |
| *Glioma* | depleted | 0.003 | 22 |
| *Melanoma* | depleted | 0.003 | 19 |
| *Non-small cell lung cancer* | depleted | 0.003 | 20 |

In many cancers, the miR-200 family of miRNA is depleted during transformation of cells from an epithelial (less motile) to a mesenchymal (more motile) cell type. An analysis of the National Cancer Institute’s 60 cell line library (NCI-60) identified a strong association between lower levels of miR-200c and a more mesenchymal cell type (36,39). Previous work from our laboratory identified a loss of α-1,2-fucosylation, as observed by the lectin UEA-I, in more mesenchymal cell types in the same cell library. We tested whether FUT1 levels correlated with miR-200c and α-1,2-fucosylation levels within the NCI-60 dataset (**Fig. 7**, **Dataset S3**). Mapping FUT1 expression across the NCI-60 shows strong correlation between miR-200c, α-1,2-fucosylation and FUT1 (*p <0.0001*).

**Figure 7.**
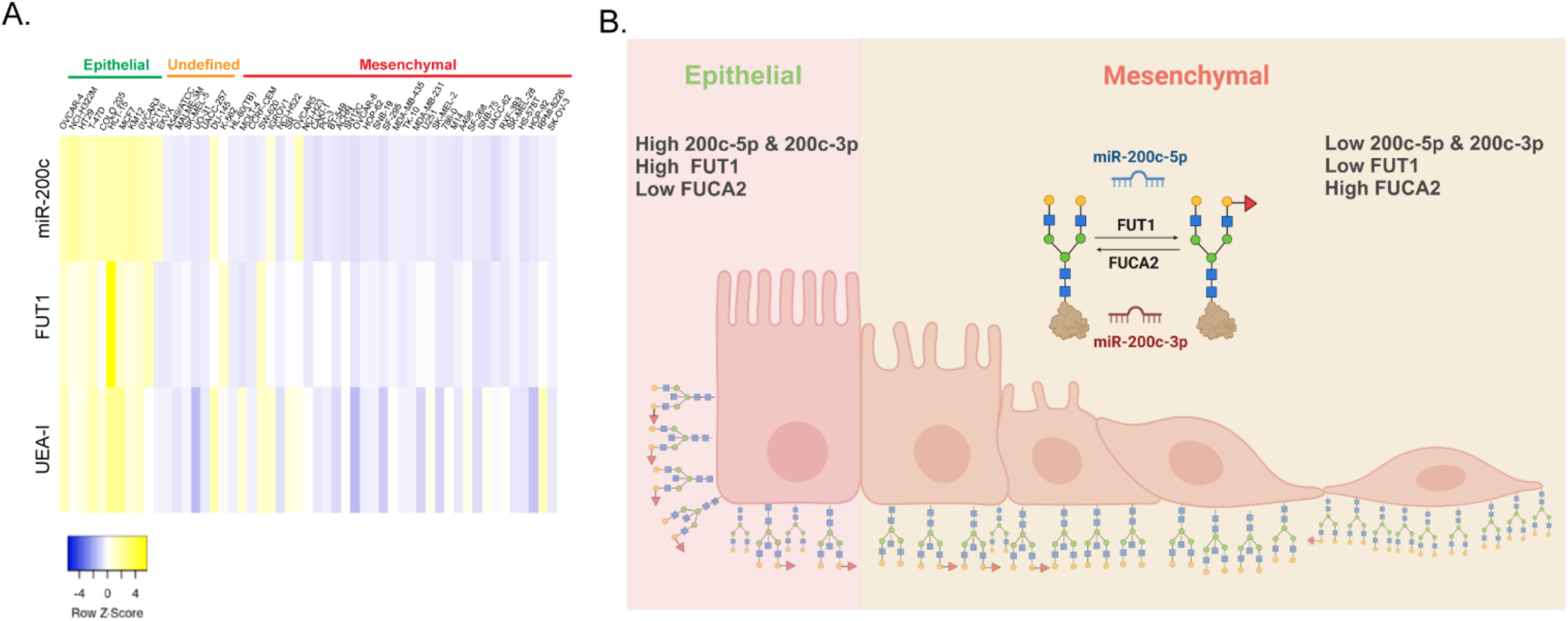
miR-200c regulates α-1,2-Fucosylation during EMT. (A)Heatmap showing the change in FUT1 expression, UEA-I binding, and miR-200c-3p change during EMT across the NCI60 library (yellow: higher expression or binding, blue: lower expression or binding. (B) Schematic representation depicting the synergistic role of miR-200c-5p and −3p in regulating FUT1 and FUCA2 during EMT. The expression of both 200c-5p and - 3p decreases during EMT, resulting loss of FUT1 upregulation by 200c-5p and FUCA2 downregulation by 200c-3p and ultimately leading to reduced α-1,2-Fucosylation in mesenchymal cells.

## Discussion

The α-1,2-fucosyltransferase FUT1 underpins blood type and has roles in establishing the gut microbiota and in cancer progression. Our high-throughput analysis of the miRNA regulatory landscape of FUT1 found bidirectional regulation with high numbers of miRNA upregulating this enzyme (80% of hits, **Fig. 2B**). We confirmed our miRFluR assay results for FUT1 in multiple cell lines of different cell types, validating that our assay identifies modulators of the endogenous FUT1 enzyme (**Figs. 3 &5**). We also observed an impact of miRNA on α-1,2-fucosylation, the end product of FUT1 enzymatic activity, with up-miRs increasing the level of the glycan epitope and down-miRs decreasing it (**Fig. 4**). Our data showcases once again that bidirectional regulation by miRNA is a common biological event.

Work from our laboratory and others has shown that upregulation by miRNA occurs through direct binding to the 3’UTR of mRNA. Although the mechanisms underlying upregulation are yet unclear, it has been shown that upregulation requires AGO2 and is independent of downregulation (13,17). Downregulation by miRNA typically requires binding in the seed region of the miRNA to the mRNA (34). In contrast, previous work from our laboratory has shown that many validated upregulatory sites found via unbiased approaches have non-canonical patterns of binding (16–21). In line with this, we identified two up-miR sites in this work (miR-200c-5p and miR-361-3p, **Fig. 6**) which were both non-canonical, with bulges on the 3’UTR side within the region of seed binding. This data adds to the growing body of evidence that upregulatory miRNA predominantly utilize a different binding modality (non-canonical) than downregulatory interactions.

In previous work we found that miRNA that target a specific gene can help identify the biological functions of that gene (miRNA Proxy Hypothesis) (12, 16, 19). This was found to be true for both up- and downregulatory miRNA interactions. Enrichment analysis of the upregulatory miRNA for FUT1 identified a dominant role for this enzyme in cancer biology. In general, miRNAs upregulating FUT1 are depleted in cancers (Table 1), consistent with previous works that found a loss of α-1,2-fucosylation is a hallmark of multiple cancers including esophageal cancer and melanoma (8,22,28,35). FUT1 has also been reported to increase host susceptibility to bacterial pathogen invasion in human and animal models (29–31). In line with this, up-miRs of FUT1 are enriched in microRNAs associated with the bacterial invasion of epithelial cells. FUT1-upregulating miRNAs were depleted in IL–17–related pathways; implying that reduced FUT1 expression may be associated with stronger pro-inflammatory IL-17 activity. Evidence for this comes from data in an IMQ-induced psoriasis model in which *FUT1*-deficient mice exhibit more severe inflammation upon IL-17a treatment, including elevated CXCL1 expression and increased neutrophil infiltration (38). Our analysis provides additional evidence for our miRNA Proxy Hypothesis, positioning miRNA as a key regulatory layer within biological networks.

In previous work, we found a strong correlation between the miR-200 family and fucose levels in cancer cells (28). A closer analysis of our data found that this association was not to general fucose, but rather was specific to α-1,2 fucosylation (**Fig. 7**). In our initial work, we identified the fucosidase FUCA2 as a target of miR-200c-3p and hypothesized that higher levels of this enzyme might result in the loss of fucosylation (28). However, our findings that miR-200c-5p is an upregulator of FUT1 provides a synergistic mechanism that may explain the strong association of α-1,2 fucosylation with expression of the miR-200c precursor from which both miR-200c-3p and −5p derive. In line with this Rohdes et al., demonstrated that both strands of another miR-200 family member, miR-200b are physiologically expressed and both the −3p and −5p strands cooperatively repress EMT through the RHOGDI pathway (40). Our findings provide evidence that the same pre-miRNA can bidirectionally regulate genes in a network to create the same biological effect through the use of different miRNA strands. Interestingly, while miR-200c-5p upregulates FUT1 and fucosylation, this same miRNA downregulates CMP-sialic acid synthetase, which synthesizes CMP-sialic acid, the nucleotide precursor for sialylation (19). In general, sialylation is highly upregulated in metastasis of many cancers, including melanoma, breast, and pancreatic cancer, where the miR-200 family is decreased (42–43). Thus, α-1,2 fucosylation and sialylation may be inversely regulated by the same miRNAs resulting in the same biological outcome. Overall, our data provides evidence that the impact of miRNA, the direction of which which is dependent on the target gene, is coordinated to control specific biology (in this case cancer transformation and metastasis).

## Experimental Procedures

### Cloning pFmiR-FUT1 sensor

FUT1 3’UTR (ENST00000310160.3, 2173 nucleotides) was amplified from cDNA (MCF7 cell line) using Q5 High-Fidelity DNA polymerase (Cat# M0492L, New England Biolabs (NEB)) according to PCR conditions provided by the company. NheI and BamHI restriction sites were used for cloning FUT1 3’UTR downstream of Cerulean in pFmiR-empty backbone using the standard NEB ligation protocol. The cloned sequence was verified by Sanger sequencing (Molecular Biology Services Unit, University of Alberta). NucleoBond Xtra Maxi EF (Ref. 740424.50, Macherey-Nagel) was used for large-scale endotoxin-free DNA preparation of the verified plasmid-pFmiR-FUT1. The plasmid map for pFmiR-FUT1 and its corresponding 3’ UTR can be found in **Figs. S1 & S2**.

### Cell Lines

Cell lines (HEK293T, HT29, and A549) were purchased directly from the American Type Culture Collection (ATCC), and A375 cells were generously supplied by Prof. Eva Hernando’s lab at New York University (NYU). HEK293T and HT29 were cultured in Dulbecco’s Modified Eagle Medium (DMEM), and A549 cells were cultured in Ham F-12K medium. Media was supplemented with 10% Fetal Bovine Serum (FBS), and cells were incubated under standard conditions (5% CO_2_, 37℃). All cells used were below passage 12.

### miRFluR High-throughput Assay

miRNA mimic library was purchased from (Dharmacon). Mimics were resuspended in ultrapure nuclease-free water (REF. #: 10977-015, Invitrogen) and aliquoted into black 384-well clear black bottom plates. In each plate, 6 replicates of Non-Targeting Control (NTC) were added, and the remaining wells were loaded with mimics at a concentration of 2 pmol. Plates were stored at −80 °C, and on the day of the experiment, they were thawed at room temperature for 15 minutes (min). 30 ng of the pFmiR-FUT1 sensor in 5 µl Opti-MEM, followed by 0.1 µL Lipofectamine™ 2000 in 5 µl Opti-MEM premixed and incubated at room temperature (RT) for 5 min, was added to the well and spun for 30sec at 300RPM. The mixture was allowed to incubate at RT in the plate for 20 min. After the incubation, 25 µL of HEK293T cells with a density of 400,000 cells/ml in phenol red-free DMEM supplemented with 10 % FBS were added to the plate and spun again 30sec at 300RPM. Plates were incubated at 37℃, 5% CO_2_. After 72 h, the fluorescence signals of Cerulean (excitation: 433 nm; emission: 475 nm) and mCherry (excitation: 587 nm; emission: 610 nm) were measured using SYNERGY H1, BioTek, Gen5 software, version 3.08.01.

### Data Analysis

Each well of each plate was scanned by a plate reader to measure the fluorescence signal of Cerulean and mCherry to calculate the ratio. Each miRNA contained three replicates of Cerulean/mCherry ratio values. The mean ratio, standard deviation (SD), and % error (100×SD/mean) were then calculated for each miRNA. At this stage, miRNAs were subjected to a quality control (QC) process where any miRNA with 15% or higher error was eliminated from the final data analysis. Additionally, if any plate had more than 50% of its total miRNA eliminated, that plate was also removed from the final data analysis. A total of 985 miRNAs passed the QC from a library of a total of 2601 miRNAs. After that, the mean ratio of each miRNA in a plate was divided by the median of the plate to calculate the normalized value. Data from all plates were then combined, and a z-score was calculated. A z-score of ±1.96 (95% confidence interval) was set as a threshold for both up- and downregulatory miRNAs. **(see Fig. 2B and Dataset S1)**

### Western Blot

Cells were seeded at a 100,000 cell/ml density in a six-well plate 24 h prior to transfection. On the day of the transfection, 50 nM miR mimic/anti-miRs in 250 μL of OptiMEM is prepared and added to a premixed 5 µL Lipofectamine reagent in 250 μL of OptiMEM. The mix was incubated at RT for 20 min. The media from the cells was removed, and fresh 1.5 ml media was added. The transfection mix was added to the cells after 20 min. Cells were lysed with RIPA buffer supplemented with protease inhibitor after 48 h, and the total protein concentration was quantified using the BCA assay. Western blot analysis was conducted for FUT1 in three cell lines. For each Western blot, an miR-8066 (NTC*) was run. Each transfection and western was repeated a minimum of three times in different passage number cells, and the average quantified result was taken.

For Western blot analysis, 50 µg of protein for HT29 and A549 and 100 µg for A375 cells was added to 4x loading buffer with β-mercaptoethanol, heated at 97 °C for 7 min, and run on a 4-15% gel (SDS-PAGE) using standard conditions. Proteins were then transferred from the gel to the Nitrocellulose membrane (nitrocellulose, Invitrogen,Cat#:IB23002) using iBlot2 Transfer Stacks and the iBlot2 transfer device (Invitrogen) using the standard protocol (P_0_). Blots were then incubated with Ponceau S Solution (Boston BioProdcuts, catalog #ST-180) for 7 min and the total protein levels were imaged using the protein gel mode (Azure 600, Azure Biosystems Inc.). Blots were blocked with 5% milk in TBST buffer (TBS buffer plus 0.1% Tween 20) for 1 h 55RPM on a rocker (LSE platform rocker, Corning) at RT. Next, blots were incubated with polyclonal Rabbit FUT1 antibody (Biosource Cat# MBS9414746) (1:200 with 5% milk in TBST) overnight at 4°C 55RPM on the rocker. After an overnight incubation at 4℃, blots were washed 1 x rinse and 4 × 3min with 0.1% TBST buffer. A secondary antibody was then added (Invitrogen Goat anti-rabbit IgG-HP Cat# 65-6120) 1: 500 in 5% milk in TBST and incubated for 1 h at room temperature with shaking 55RPM. Blots were then washed 4 × 2min with 0.1% TBST buffer. Blots were developed using Clarity and Clarity Max Western ECL substrate according to the manufacturer’s instructions (Bio-Rad Cat# 1705061). Membranes were imaged in chemiluminescent mode (Azure 600, Azure Biosystems Inc).

### Data analysis of Western blot

All analyses were done for a minimum of biological triplicate. Images were quantified using Image J software .tiff files were used for analysis (Image J 1.54g, Java 1.8.0_345). For each lane, the signal was normalized to the Ponceau for that lane. For each blot, the Ponceau normalized signal for miRNA mimics was divided by the NTC* to give the normalized signal shown in all graphs. We tested for statistical significance using a one-sample *t*-test. **(see Table S4)**

### UEA-I staining

Fluorescein (FITC) conjugated *Ulex Europaeus Agglutinin I* (UEA-I) lectin was purchased from Thermofisher Cat# L32476). Sterile 22×22 no. 1 coverslips were placed on a 35 mm plate. Cells were seeded at 50,000 cells/ ml and treated with miRNA mimics/ anti-miRs as described in the Western Blot section. 48 h post-transfection, cells were washed with 1 ml TBS containing 0.1% BSA and then fixed with 4 % paraformaldehyde at room temperature for 10 min. After fixation, cells were washed with sterile TBS containing 0.1% BSA buffer (3 x 3 min) and blocked with 1% BSA in TBS for 1h in the incubator (37^°^ C, 5% CO_2_). FITC-UEA-I was then added to the cells at a dilution of 1:500 in TBS. Cells were incubated at RT in the dark. After 1 h of incubation, cells were again washed with TBS containing 0.1% BSA (3x) and counterstained with Hoechst 33342 (Invitrogen Cat# H1399) at room temperature for 2 min in the dark. The coverslips were then mounted onto slides with mounting media (90% glycerol in TBS) and imaged with a Zeiss fluorescent microscope (Camera: Axiocam 305 mono, software: ZEN 3.2 pro). All analysis was done in biological triplicate.

To confirm the specificity of *Ulex europaeus agglutinin I* (UEA-I) lectin staining, a competitive inhibition control was performed using free L-fucose. Cells were incubated with FITC-conjugated UEA-I in the presence or absence of 100 mM L-fucose for 30 min at room temperature. L-fucose competes with α-1,2-fucosylated glycans for UEA-I binding and therefore serves as a negative control for lectin specificity. After incubation, cells were washed with TBS containing 0.1% BSA (3x) and nuclei were counterstained with Hoechst 33342 (Invitrogen Cat# H1399). Fluorescence images were acquired using identical microscope settings for all conditions **(Fig. S7).**

### Flow Cytometry

Cells well were seeded (100,000 cells/ml) into a 6-well plate and incubated under standard conditions (37° C, 5% CO_2_, media). After 24 h, cells were transfected with miR mimics or anti-miRs as previously described. At 24 h post-transfection, the transfection media was removed and replaced with fresh media. After an additional 48 h, cells were washed with 1 ml sterile wash buffer (TBS, 1 % BSA) 3 x and then trypsinized (200 µl / well, 0.25 % trypsin). The detached cells were diluted with wash buffer and centrifuged at 300 x g for 7 min. Cell pellets were then resuspended in ice-cold wash buffer, the cell number was adjusted to 50,000 cells/ml for each condition, and cells were pelleted. Cells were resuspended in 20 µg/ ml UEA-I lectin in 100 µL TBS, 0.1% BSA, and incubated in the dark for 30 min. Cells were centrifuged at 300 x g for 7 min and subsequently washed 3 x. Following the third wash, cells were resuspended in 800 μL FACS buffer (TBS,1 % BSA, 10 % FBS) and analyzed with an Attune NxT Acoustic Focusing cytometer. Cells were gated using forward and side scatter to ensure only live cells were analyzed.

To validate the specificity of UEA-I binding in flow cytometry, a competitive inhibition assay was performed using free L-fucose. Cells were incubated with FITC-conjugated UEA-I in the presence or absence of 200 mM L-fucose for 30 min at 4 °C. L-fucose competes with α-1,2-fucosylated glycans for UEA-I binding and therefore serves as a specificity control. After incubation, cells were washed with PBS and analyzed by flow cytometry. Mean fluorescence intensity (MFI) was calculated to quantify UEA-I binding **(Fig. S8)**.

### Multi-Site Mutagenesis on pFmiR:FUT1

miRNA binding sites of FUT1 3’UTR for miR-200c-5p, miR-361-3p, and miR-29c-5p were predicted using RNAhybrid (37). The first predicted site with the lowest delta G value was mutated to the corresponding miRNA sequence using Q5^®^ Site-Directed Mutagenesis kit according to the manufacturer’s instructions. Primers for this experiment were designed using the NEB base changer tool and purchased from Integrated DNA Technologies (IDT). The mutation’s incorporation was confirmed by Sanger and nanopore sequencing and maxi-prepped for further miRFluR analysis.

### siRNA Knock Down of FUT1

ON-TARGETplus siRNA reagents against FUT1 in a smart pool format and ON-TARGETplus Non-Targeting Control Pool (NTP) were purchased from Dharmacon (Horizon Discovery, CA). A375 cells were seeded in six-well plates (100,000 cells/well) and cultured for 24 h in appropriate media. Cells were then transfected with each of the siRNA pools (25, 50, 75 nM si-FUT1 NTP (50nM NTP), Dharmacon, Horizon Discovery) using Lipofectamine™ RNAiMAX transfection reagent (catalog #: 13778150, Thermofisher) following the manufacturer’s instructions. After 48 h, cells were lysed using RIPA Lysis buffer supplemented with protease inhibitors (Thermofisher, catalog #: 89900), and lysates were quantified using BCA assay (Micro BCA™ Protein Assay Kit, catalog #: 23235) and were analyzed for Western blot as previously described. Blots were incubated with antibodies and processed as described before. (**see Fig. S6**)

## Data availability

The authors declare that all data can be found in this document and its supporting files.

## Supporting information

This article contains supporting information:

Figs. S1-S8

Tables S1-S5

Datasets S1 (separate Excel files)

## Acknowledgements

We thank Prof. Eva Hernando for generously providing A375 cells.

## Author contributions

Conceptualization: T.T.B., L.K.M.; Methodology: T.T.B., C.T., L.K.M.; Investigation: T.T.B., Validation: T.T.B.; Formal Analysis: T.T.B., L.K.M.; Visualization: T.T.B., L.K.M.; Writing-Original Draft: T.T.B., L.K.M; Supervision: L.K.M.

## Funding and additional information

Funding for L.K.M. comes from the Canada Excellence Research Chairs Program (CERC in Glycomics). Flow cytometry experiments were performed at the University of Alberta Faculty of Medicine & Dentistry Flow Cytometry Facility, RRID:SCR_019195, which receives financial support from the Faculty of Medicine & Dentistry and Canada Foundation for Innovation (CFI) awards to contributing investigators.

## Abbreviations

The abbreviation used are as follows:

FUT1: Fucosyltransferase
miRNA: microRNA
AGO2: Argonaute 2
3’UTR: 3’-untranslated regions
mRNA: messenger ribonucleic acid
siRNA: small interfering RNA
NTP: non-targeting control pool
NTC: non-targeting control
down-miR: downregulatory miRNA
up-miR: upregulatory miRNA
HRP: horseradish peroxidase
FITC: fluorescein isothiocyanate
UEA-I: Ulex Europaeus Agglutinin I
kDa: kilodalton
WT: wild-type
MUT: mutant
IgG: immunoglobulin G
SDS-PAGE: sodium dodecyl sulfate polyacrylamide gel electrophoresis
PCR: polymerase chain reaction

