## Supplemental Information for "microRNAs bidirectionally regulate FUT1 to modulate α-1,2-fucosylation and cancer-associated biology"

**This file contains:**

**Figures: S1-S8**

**Table: S1**

**Dataset: 1-3**

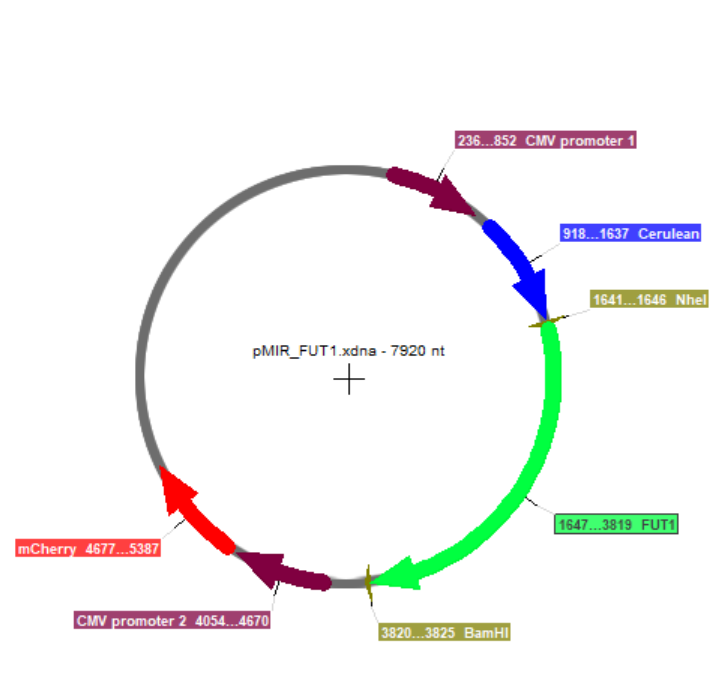

**Figure S1. pFmiR-FUT1 map**  pFmiR-FUT1 plasmid map. (Created using SnapGene)

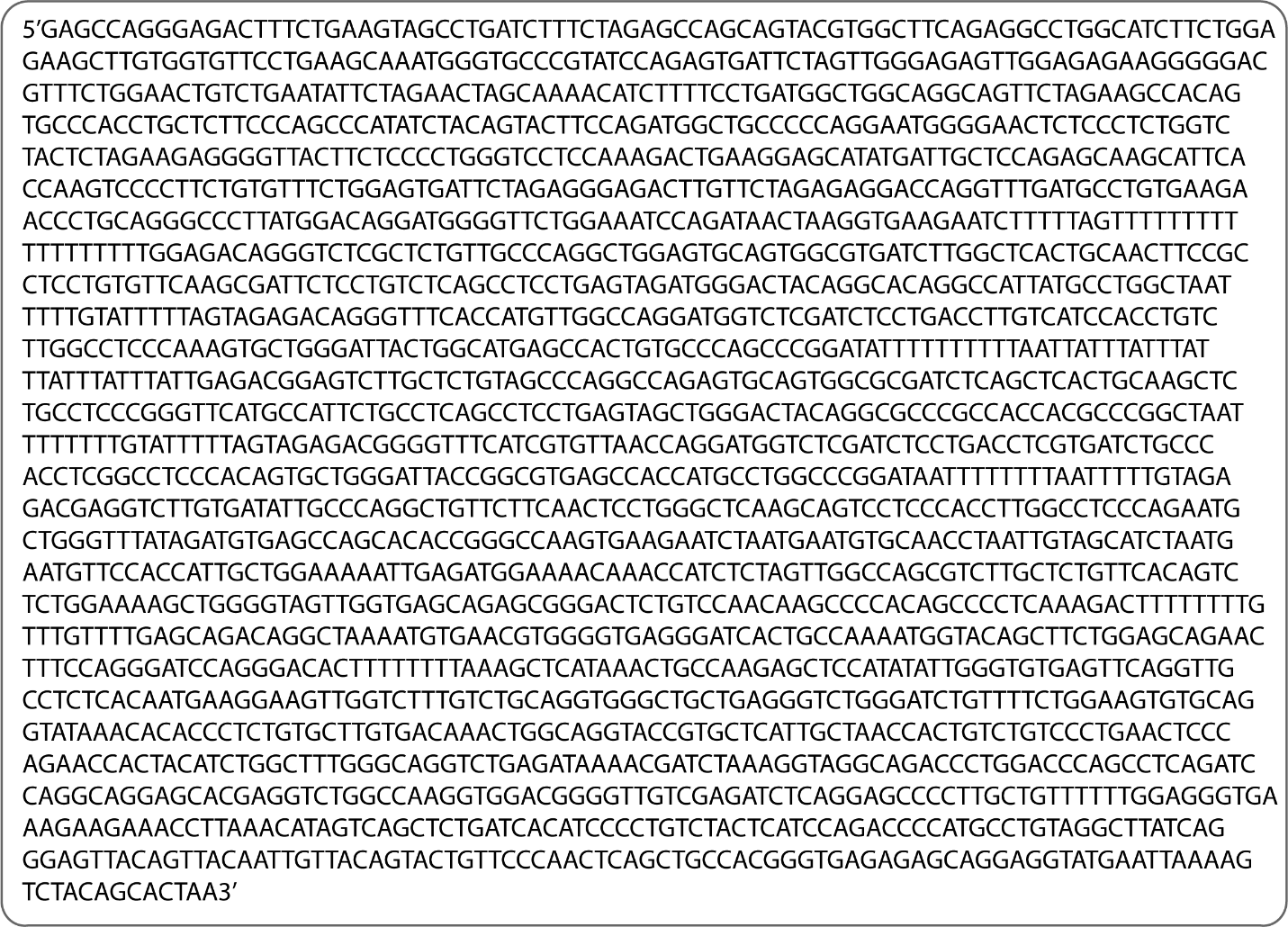

**Figure S2. FUT1 3’UTR sequence.** FUT1 3'UTR sequence (2173 base pairs).

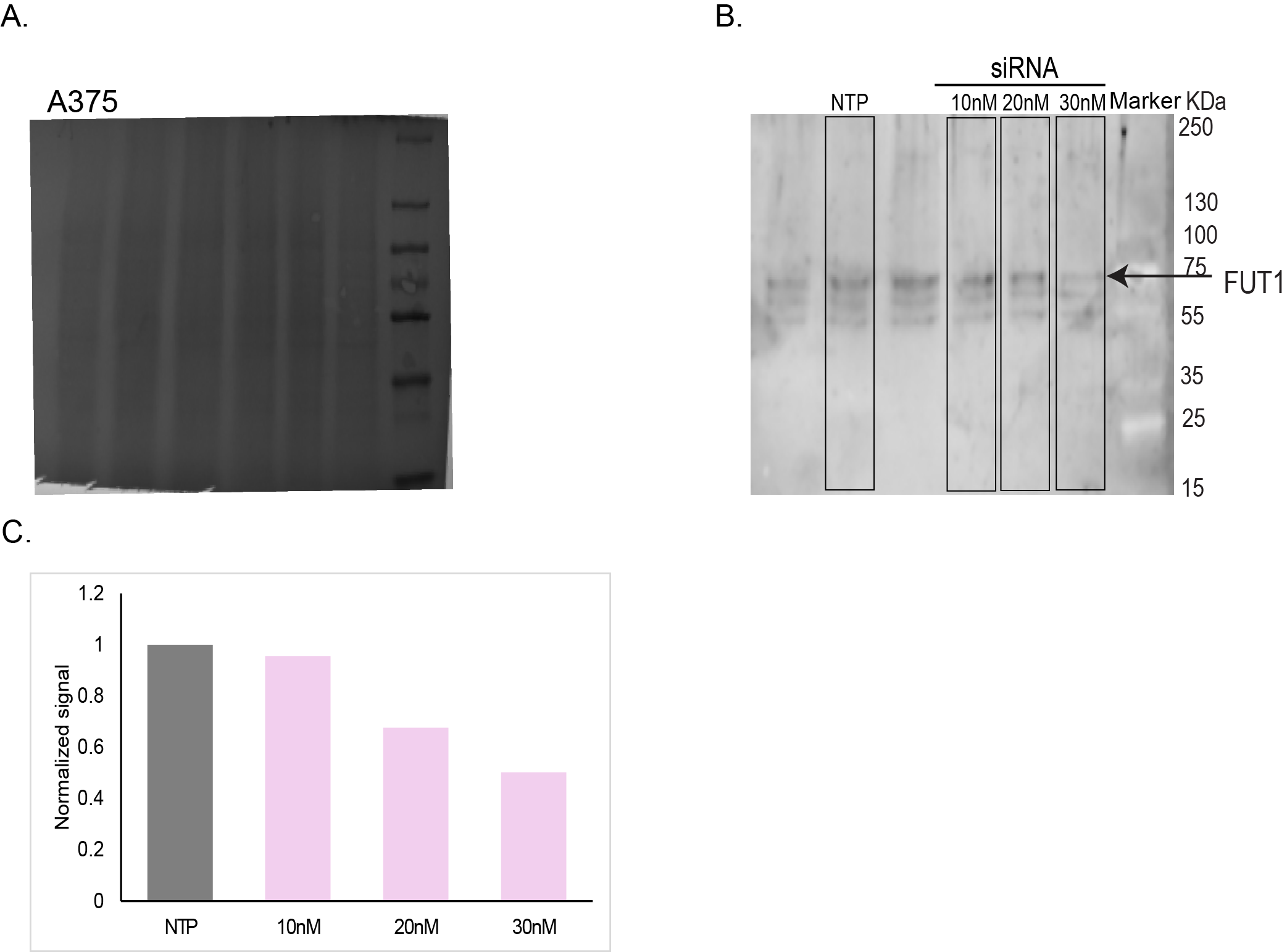

**Figure S3. Validation of ant-FUT1 antibody.**  siRNA against FUT1 was transfected into A375 using the standard protocol, NTP (30nM), siRNA(10,20,30nM). A. Ponceau staining. B. Whole western blot approx. 70kDa validated FUT1 band. C. Western blot quantification of the top band.

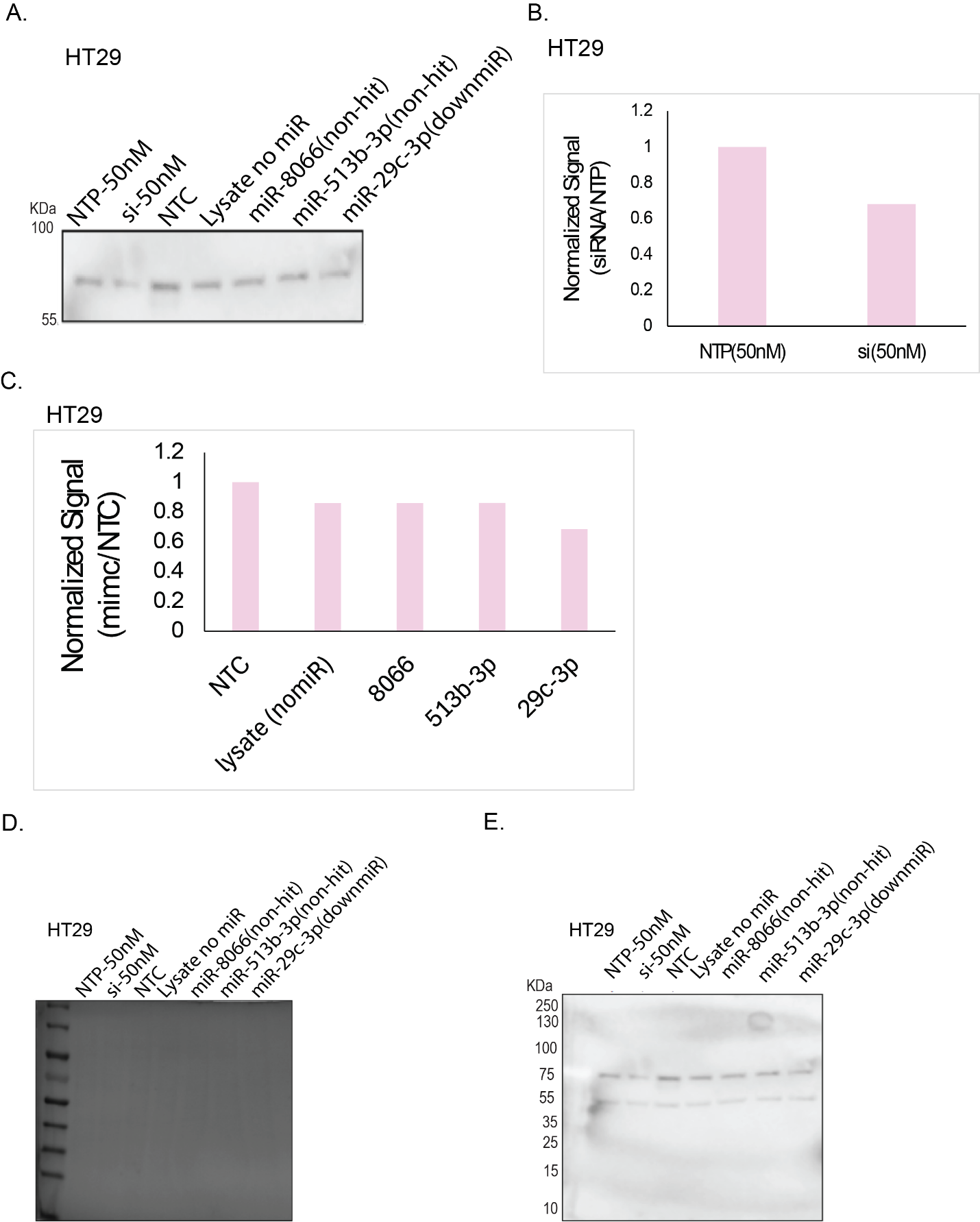

**Figure S4. Identification of a new non-targeting miRNA control (NTC*).**
Candidate miRNAs from the **median range of the miRFluR dataset** were screened for their effect on FUT1 expression. (A) Western blot showing FUT1 levels following transfection with candidate miRNAs. (B) Quantification of normalized FUT1 signal comparing non-targeting pool (NTP) and siRNA controls. (C) Comparison of normalized FUT1 expression across tested candidates and miRNA controls. (D) Ponceau staining confirming equal loading. (E) Full western blot. **miR-8066** produced a signal comparable to the lysate-only control, indicating no measurable effect on FUT1 expression, and was therefore selected as the **non-targeting control (NTC*)** for subsequent experiments.

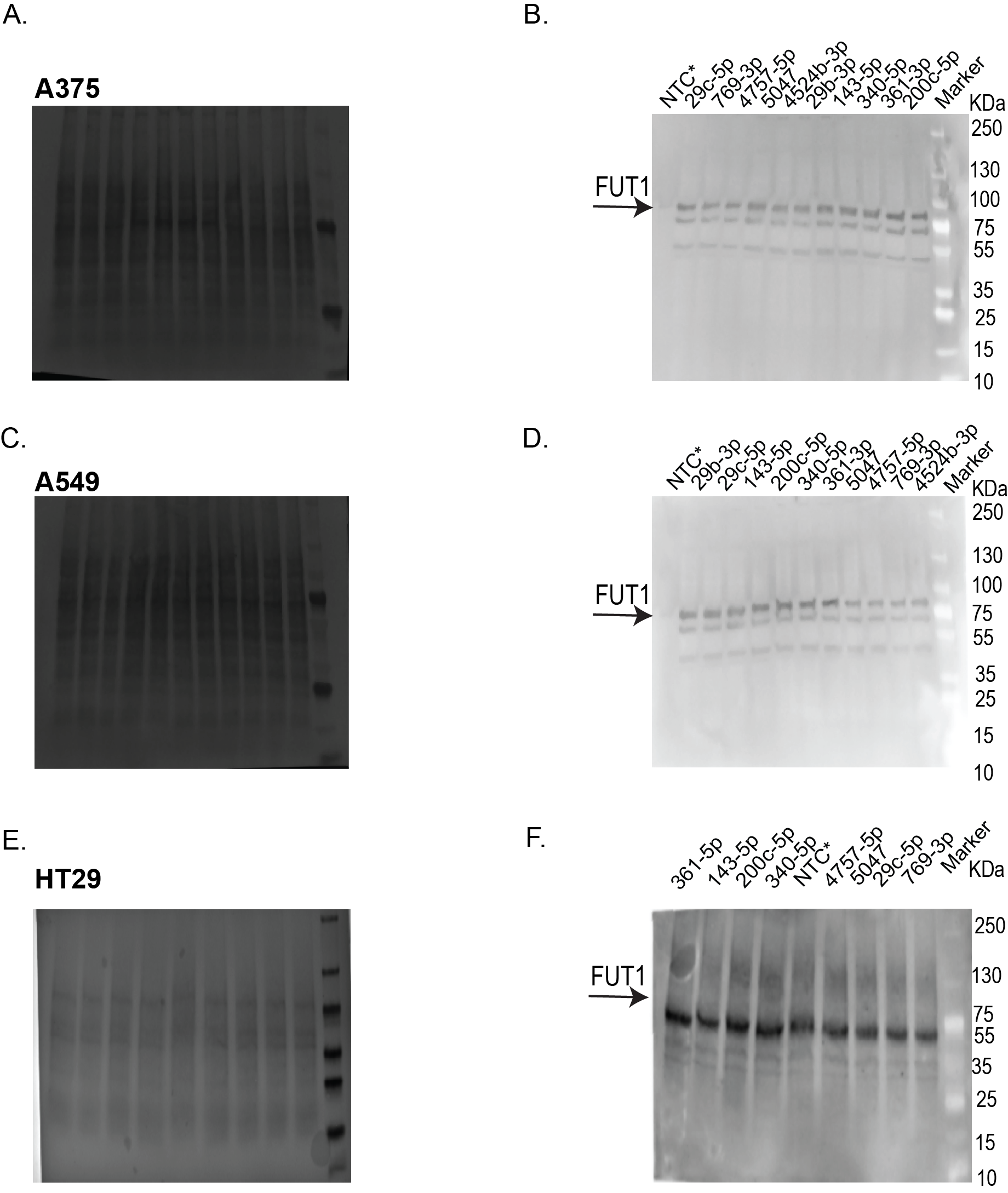

**Figure S5.** **Ponceau and whole Western blots for data shown in Fig. 3A.**  (A) Ponceau staining of blots used in Fig. 3A top (A375), (B) Whole Western blot for data shown in Fig. 3B (A375), (C) Ponceau staining of blots used in Fig. 3A middle (A549), (D) Whole Western blot for data shown in Fig. 3C (A549), (E) Ponceau staining of blots used in Fig. 3A bottom (HT29), (F) Whole Western blot for data shown in Fig. 3D (HT29).

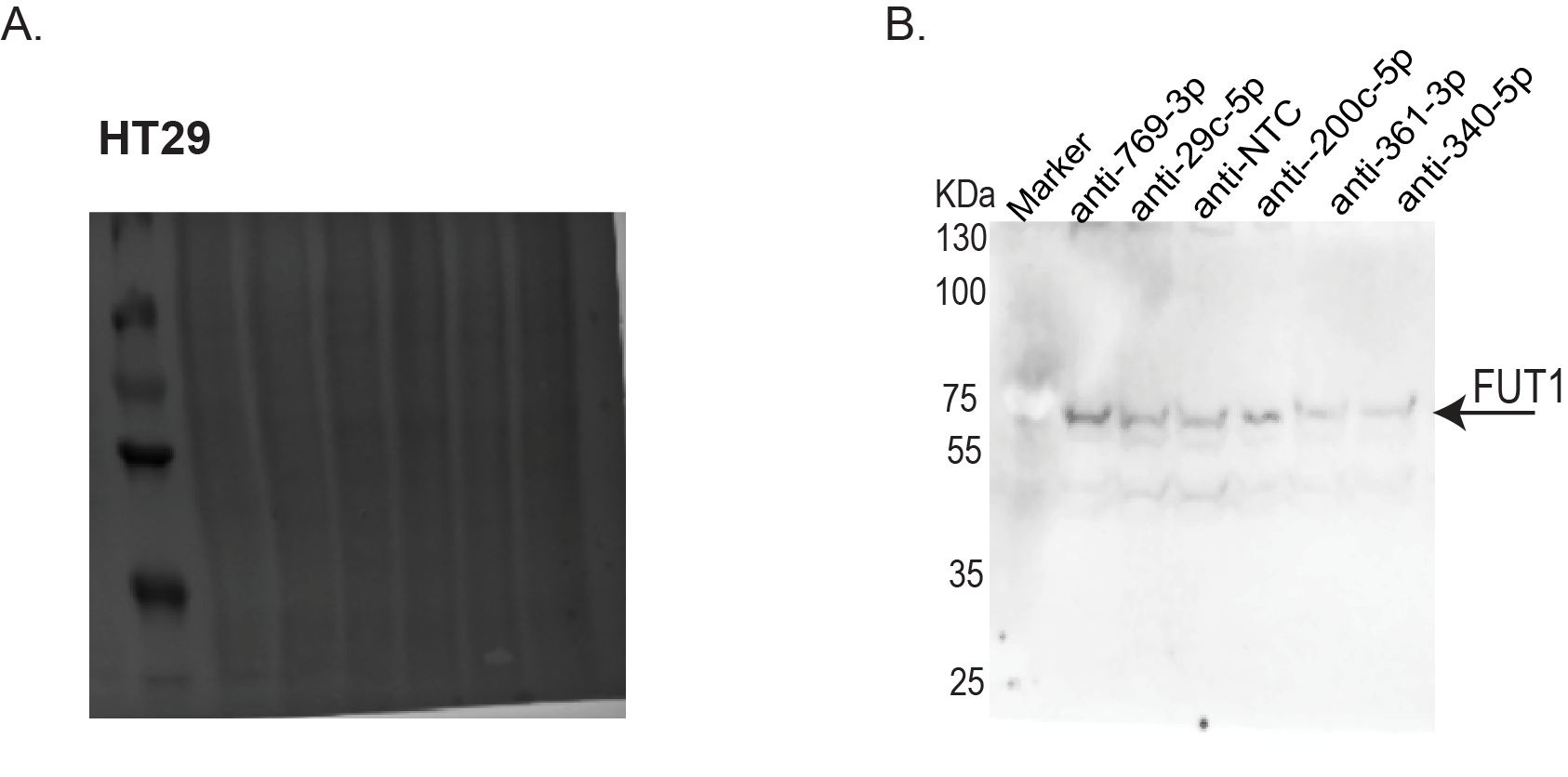

**Figure S6.** **Ponceau and whole Western blots for data shown in Fig. 5B.**  (A) Ponceau staining of blots used in Fig. 5B (HT29), (B) Whole Western blot for data shown in Fig. 5B (HT29).

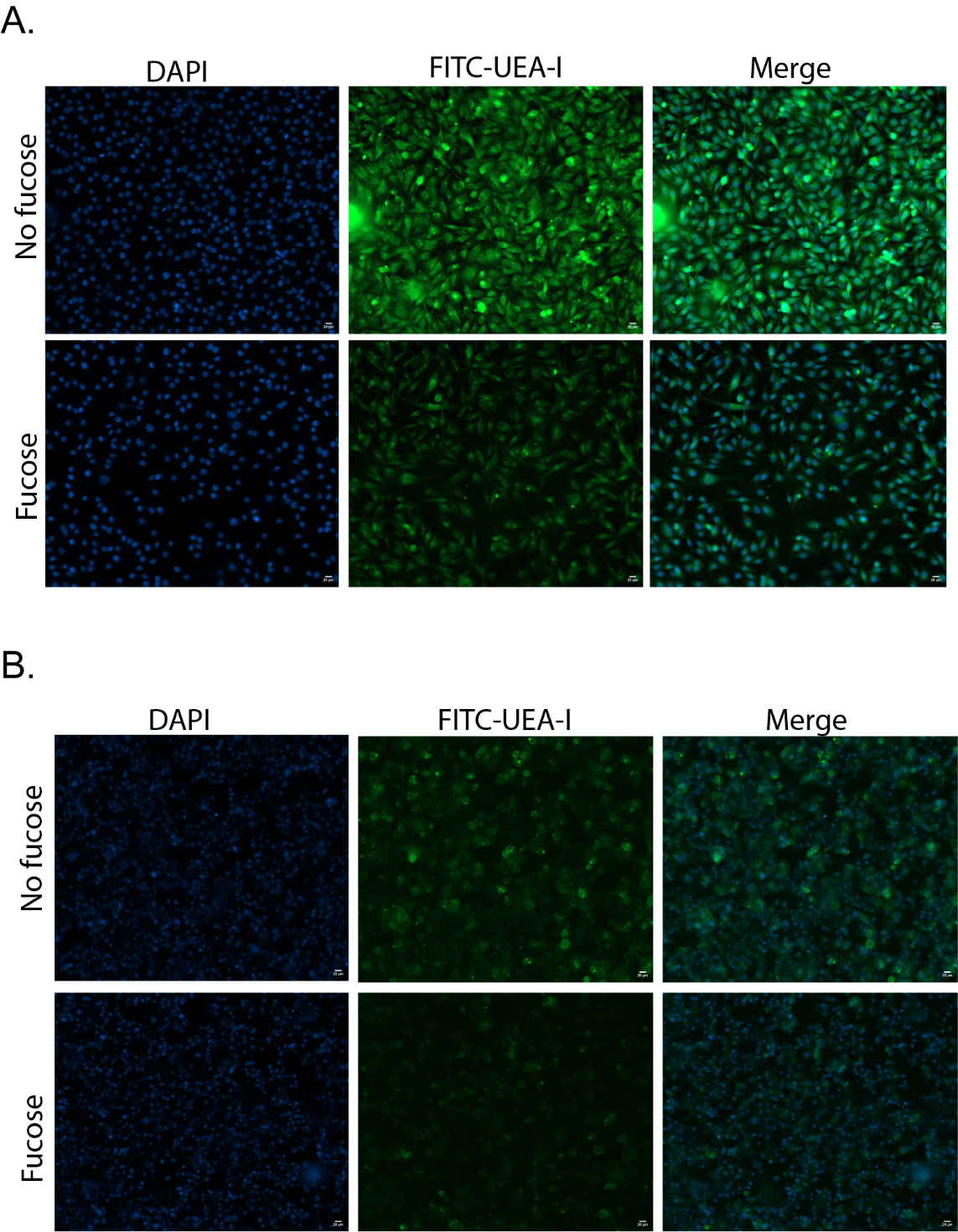

**Figure S7. Negative control for UEA-I binding lectin staining.** (A) A375, (B) HT29

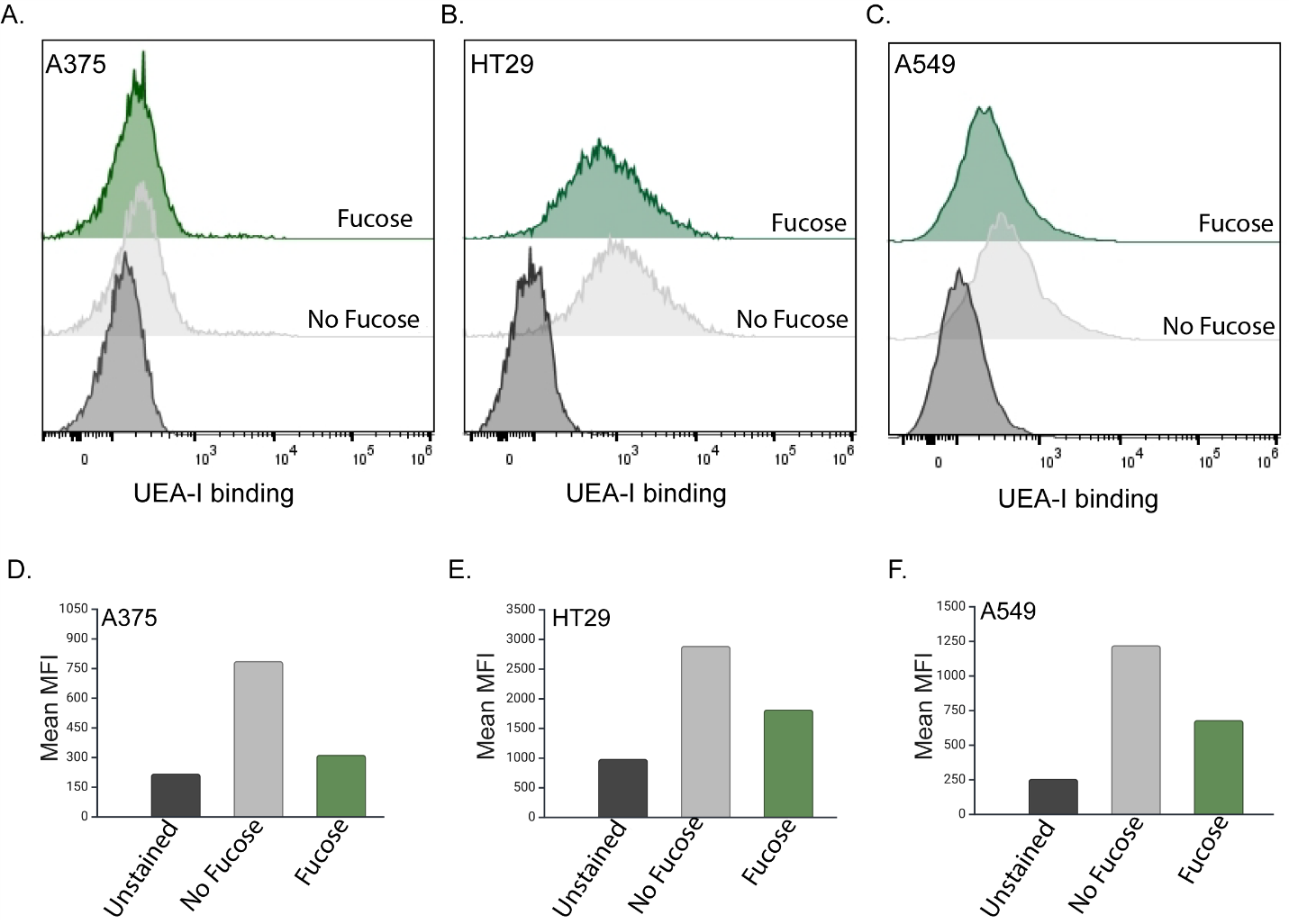

**Figure S8. Negative control for UEA-I binding flow cytometry.** (A) A375, (B) HT29

**Table S1** Statistical significance for western blot experiments using one-sample *t-*test

| **miRs/anti-miRs** | **Cell line** | | |
| --- | --- | --- | --- |
|  | **A549** | **A375** | **HT29** |
|  | **One-sample *t*-test** | | |
| Down-miRs |  |  |  |
| 29c-5p | 0.0335 | 0.0119 | 0.0043 |
| 769-3p | 0.0127 | 0.0064 | 0.0609(NS) |
| 4757-5p | 0.0192 | 0.9701 (NS) | 0.2666(NS) |
| 5047 | 0.0761 (NS) | 0.0032 | 0.1466(NS) |
| 4524b-3p | 0.0815 (NS) | 0.0089 |  |
| Up-mirs |  |  |  |
| 29b-3p | 0.1487 (NS) | 0.0085 |  |
| 143-3p | 0.0355 | 0.0121 | 0.0002 |
| 340-5p | 0.0193 | 0.0015 | 0.0038 |
| 361-3p | 0.0064 | 0.0034 | 0.0008 |
| 200c-5p | 0.0384 | 0.0074 | 0.0257 |
| Down-miRs |  |  |  |
| anti-29c-3p |  |  | 0.0063 |
| anti-769-3p |  |  | 0.0075 |
| Up-mirs |  |  |  |
| anti-200c-5p |  |  | 0.3739(NS) |
| anti-340-5p |  |  | 0.0297 |
| anti-361-5p |  |  | 0.0119 |

**Dataset S1:** Tab 1: Total dataset, Tab 2: Dataset after removing plates with high error

Tab 3: QCed data (<15% error of measurement), Tab 4: Z-scored data.

**Dataset S2:** NCI-60 library miR-200c, FUT1, UEAI expression**.**

**Dataset S3:** miEAA enrichment analysis for FUT1 upmiRs (KEGG, MNDR)
